# Structural characterization of SLYM - a 4^th^ meningeal membrane

**DOI:** 10.1101/2023.10.20.563351

**Authors:** Virginia Plá, Styliani Bitsika, Michael Giannetto, Antonio Ladron-de-Guevara, Daniel Gahn-Martinez, Yuki Mori, Maiken Nedergaard, Kjeld Møllgård

## Abstract

Traditionally, the meninges are described as 3 distinct layers, dura, arachnoid and pia. Yet, the classification of the connective meningeal membranes surrounding the brain is based on postmortem macroscopic examination. Ultrastructural and single cell transcriptome analyses have documented that the 3 meningeal layers can be subdivided into several distinct layers based on cellular characteristics. We here re-examined the existence of a 4^th^ meningeal membrane, <u>S</u>ubarachnoid <u>Ly</u>mphatic-like <u>M</u>embrane or SLYM in Prox1-eGFP reporter mice. Imaging of freshly resected whole brains showed that SLYM covers the entire brain and brain stem and forms a roof shielding the subarachnoid cerebrospinal fluid (CSF)-filled cisterns and the pia-adjacent vasculature. Thus, SLYM is strategically positioned to facilitate periarterial influx of freshly produced CSF and thereby support unidirectional glymphatic CSF transport. Histological analysis showed that, in spinal cord and parts of dorsal cortex, SLYM fused with the arachnoid barrier layer, while in the basal brain stem typically formed a 1-3 cell layered membrane subdividing the subarachnoid space into two compartments. However, great care should be taken when interpreting the organization of the delicate leptomeningeal membranes in tissue sections. We show that hyperosmotic fixatives dehydrate the tissue with the risk of shrinkage and dislocation of these fragile membranes in postmortem preparations.

## Introduction

Numerous studies have over the past decade shown that glymphatic fluid flow removes amyloid-β and other waste products of neural metabolism from brain parenchyma (1). These findings, along with the discovery of the meningeal lymphatic vessels, have sparked a new interest in brain fluid transport (2). Parallel investigations documented that glymphatic/lymphatic clearance is suppressed in aging and that the decline in brain fluid transport is accentuated in patients suffering from neurodegenerative diseases compared with age-matched non-demented control subjects (3-7). An emerging question is whether therapies that restore glymphatic/lymphatic clearance can delay the onset of dementing diseases or modify their progression (8).

One aspect of brain fluid transport that has received little attention is the functional organization of the large CSF-filled subarachnoid space (SAS) surrounding the brain. SAS is logically central for the organization of brain fluid transport, yet essentially no information exists on CSF dynamics in the SAS. Most data regarding the structure of SAS is based on histological analysis. Postmortem histological analysis of the SAS poses a potential problem since death and formaldehyde-based fixation eliminate the fluid filled spaces in both peripheral tissues (9) and brain (10, 11). We recently described the existence of a 4^th^ meningeal membrane, the <u>S</u>ubarachnoid <u>Ly</u>mphatic-like <u>M</u>embrane, SLYM, surrounding the mouse and human brain (12). SLYM was identified by the expression of Prospero Homeobox 1 (Prox1), which is a transcription factor that drives lymphatic endothelial fate (13). Consequently, Prox1-eGFP reporter mice are often used along with other lymphatic markers to identify lymphatic or lymphatic-like tissues (14). Structurally, SLYM is composed of a single cell layer that only expresses a subset of lymphatic markers, Prox1 and podoplanin, but not VEGFRC or LYVE1 (15). SLYM is therefore best characterized as a lymphatic-like meningeal layer. Structurally and phenotypically, SLYM was shown to differ from the 3 traditional meningeal membranes—including dura, the arachnoid barrier cell (ABC) layer, and pia—as well as from the arachnoid trabecula and lymphatic vessels (12). Functional in vivo analysis showed that SLYM compartmentalizes the SAS into two separate CSF pools (12), thus directing CSF transport in the SAS.

CSF is, at least in part, produced by the choroid plexus and exits the ventricles via the foramen of Magendie and Luschka (16). From here, the freshly produced CSF enters the basal cisterns, initially flowing into the cisterna magna, which is continuous with the interpeduncular cistern surrounding the circle of Willis. The interpeduncular cistern serves as a pool of CSF that is transported up along the perivascular space surrounding the 3 major cerebral arteries, the anterior, middle and posterior cerebral arteries in both rodents and human brain (1, 17). We showed that SLYM creates a roof over the pia-adjacent arteries as they follow the brain surface, thus facilitating that the freshly produced CSF in the basal cisterns is transported along the 3 major cerebral vessels (anterior, middle and posterior) rather than mixing with CSF in the larger SAS. However, online comments raised criticism of our description of SLYM. One set of critiques stated that our finding contradicted prior histological studies showing that SAS is a single large compartment. Another point of critique was that the immunolabeling analysis did not sufficiently document the ABC layer and SLYM as separate layers. The last set of critiques stated that our *in vivo* tracer injection failed to document the existence of 2 distinct SAS compartments. It was suggested that the injections created an artificial space and that one of the CSF-filled compartments was the subdural space. The subdural space —which does not exist during physiological conditions— is a compartment created by an intradural separation between the dura *per se* and the dural border layer. Yet, the same authors’ recent studies dissecting the meningeal layers provided histological and single cell transcriptome evidence for the existence of a Prox1-positive meningeal layer located below the ABC layer (18, 19), as we originally reported (12). In vivo studies also confirmed that SLYM and dura are separated by a CSF-filled compartment, but the authors concluded that it was an artificial space without explanation. Based on the histological and transcriptome analysis, the SLYM layer was labeled BFB2-3 (18, 19).

In this report, we have extended the characterization of SLYM and confirmed that SLYM is indeed a distinct membrane that surrounds the entire brain. More specifically, SLYM forms a roof covering the CSF-filled cisterns and pial perivascular spaces creating a continuous CSF-filled compartment. We provide evidence showing that SLYM immunophenotypically differs from the ABC layer and a critical literature review documents that compartmentalization of SAS, by either a single-or double-layered membrane, has been reported earlier (20, 21). In the older literature, SLYM has been described as the inner or reticular arachnoid membrane. We conclude that SLYM *in vivo* represents a distinct meningeal membrane that can fuse during fixation with the ABC layer or the pial membrane depending on the location. However, in the basal part of the brain SLYM creates two separate CSF-containing SAS compartments, even seen in paraffin sections. We speculate that this arrangement facilitates that fresh CSF produced in the ventricles is guided directly up along the cerebral arteries instead of being mixed with the larger pool of CSF surrounding the brain.

## Results

### Fixation-induced shrinkage of the brain

In the classical electron microscopy (EM) literature, SLYM is recognized as the inner arachnoid layer or the reticular layer of the arachnoid membrane. SLYM will in EM typically adhere to the ABC layer. Could the cohesion be an artificial product of displacement of the fragile meningeal membrane when preparing histological samples? To illustrate the significant shrinkage of the brain and the expansion of the subarachnoid space upon perfusion fixation, we first visualized the live and dead mouse brain *in situ* by high field magnetic resonance imaging (MRI) (**Fig. 1A**). The MR imaging clearly illustrates that, in a live young mouse, the brain fills most of the cranial cavity; and how small a volume the subarachnoid space surrounding the brain surfaces occupies. After death, the brain expands but swelling is restricted by the skull, resulting in compression of the subarachnoid and ventricular spaces (**Fig. 1A**). Our previous studies have documented that influx of CSF via the glymphatic system accounts for most of the acute edema formation after cardiac arrest (22). In sharp contrast, the perfusion-fixated brain imaged in thin paraffin sections exhibited a significant degree of shrinkage resulting in displacement of the brain within the skull cavity (**Fig. 1B-C**). The properties of skull’s bony structure do not change when exposed to fixative, in contrast to the soft tissue including the brain and the meningeal membranes. The striking reduction in brain volume during histological preparation results in the creation of large empty spaces surrounding the brain in the perfusion-fixated brain (9, 11). It appears that the membranes fuse to different surfaces during perfusion-fixation, with the ABC layer localized to the skull (stained for Cld-11, Fig. 1D, top), and the SLYM localized to the brain surface (stained for GFP, Fig1D, bottom). This further suggests the SLYM membrane is not always fused with the ABC layer and we conclude that the fragile leptomeningeal membranes are prone to displacement and fixation artifacts in histological sections.

**Fig. 1.**
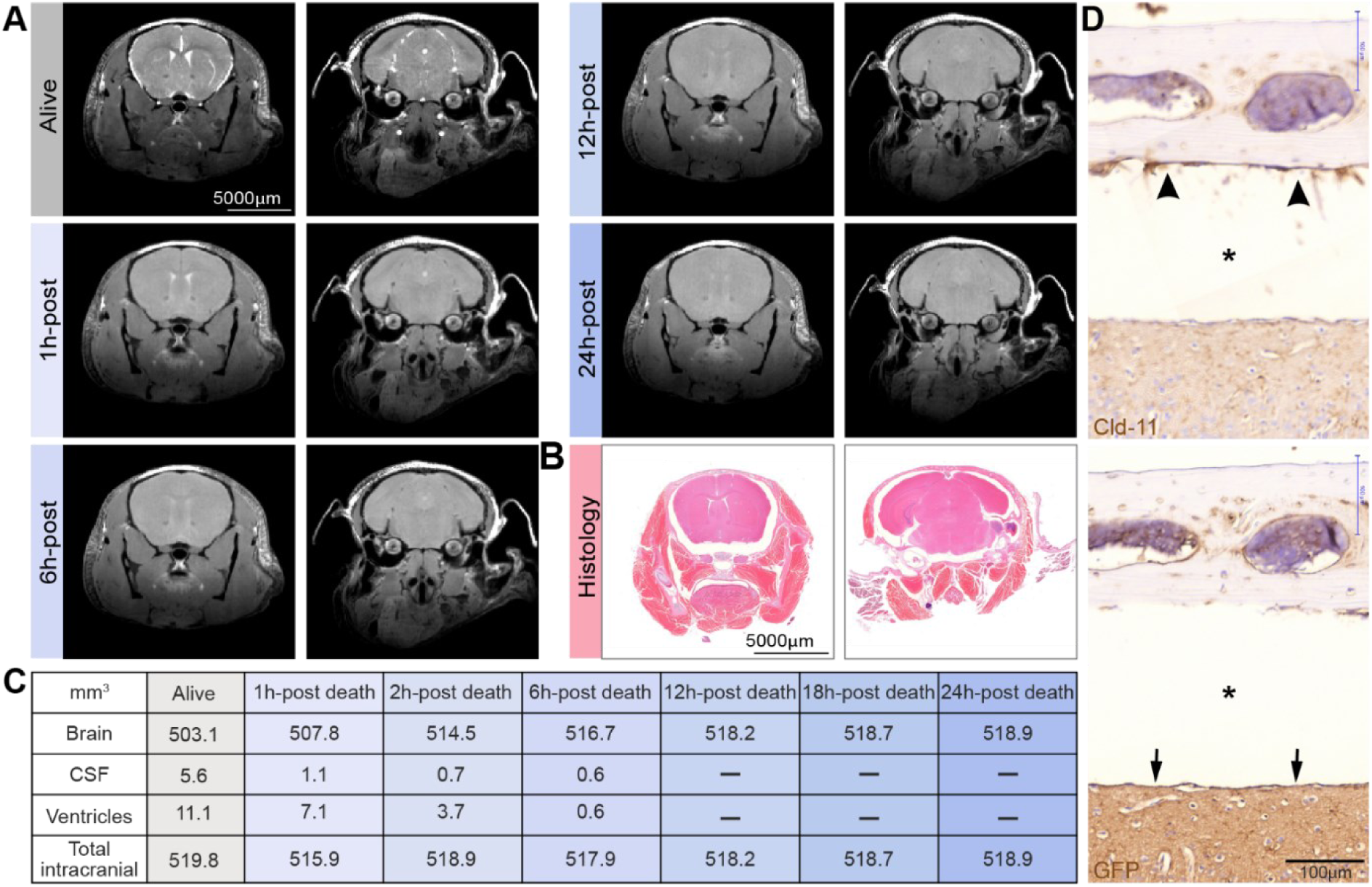
Preparation of histological sections is linked to a marked shrinkage of the brain’s fluid-filled spaces. **(A)** Magnetic resonance imaging (MRI) of a young C57BL/6JRj mouse shows that CSF in the subarachnoid space and the ventricles in a live isoflurane anesthetized mouse gradually disappear after death, indicative of brain swelling. After the live imaging session, the anesthetized mouse was killed by 100% nitrogen inhalation via a nose cone on an MR-compatible stereotactic holder, and scanning was repeated at 1, 2, 6, 12, 18, and 24 h after cardiorespiratory arrest. (**B)** Histological section of the head of a young C57BL/6JRj mouse after decalcification and preparation of parafin. **(C)** Quantification of the volume of the brain, the subarachnoid space, the ventricles and total intracranial volume based on the images displayed in panel A. **(D)** Immunohistochemical analysis of arachnoid barrier cell layer (top, arrowheads) stained for Cld-11 and SLYM stained for GFP (lower, arrows) in two consecutive sections prepared from a Prox1-eGFP mouse. Note the considerable shrinkage of the brain (asterisk) in the sections.

### SLYM creates a unique CSF compartment encompassing the cisterns and pial perivascular spaces

To interrogate the functional significance of SLYM covering the brain, Prox1-eGFP mice were injected with either a small size tracer (Evans Blue, 0.9 kDa) or a larger tracer—Bovine Serum Albumin—coupled to a fluorophore (BSA-Alexa Fluor 647, 67 kDa). The tracers were injected via a cannula placed in the cisterna magna and allowed to circulate for 30 min in anesthetized mice (ketamine/xylazine, K/X) before harvest (**Fig. 2A**). CSF tracers delivered by this procedure are initially transported into the basal cisterns followed by an influx along the perivascular space surrounding the large cerebral arteries and driven by arterial pulsations (10). The mice were intracardially perfused with phosphate-buffered saline (PBS) to eliminate any signal from the vascular compartment. First, the unfixed brains were rapidly extracted and imaged using a fluorescent macroscope to avoid fixation artifacts (**Fig. 2B-C and Sup. Figs. 1 and 2**) (11). Macroscopic imaging of the intact whole brains allowed the visualization of Evans Blue and BSA-647 confined within the inner CSF-filled SAS compartment below the SLYM layer. Visual inspection revealed that both the small and large tracers were confined within the cisterns including cisterna magna, the pontine cistern, the interpeduncular cistern, the cerebellopontine angle cistern, and several other minor cisterns. The perivascular spaces surrounding the anterior, middle, and posterior cerebral arteries were also clearly delineated by both CSF tracers (**Fig. 2B-C and Sup. Fig. 1B**). Close-up images showed the perivascular location of the tracer and Prox1-eGFP+ cells covering the tracer-filled periarterial spaces creating one continuous, closed CSF-filled SAS compartment (**Fig. 2C**). High-magnification images were combined with whole-mount confocal and two-photon imaging to further investigate whether the tracers were contained below the SLYM layer. When extracting the brains from Prox1-eGFP reporter mice, we noted that SLYM adheres to the surface of the brain rather than to dura, which stays attached to the skull (**Fig. 2D**) (23). In contrast, the ABC layer detected in the fixed skull immunolabeled positive for dipeptidyl peptidase IV (DPP4). an enzyme suggested to participate in neuropeptide metabolism at the outer arachnoid, adhered to dura (**Fig. 2D**) (19). These observations show that SLYM creates a unique CSF compartment encompassing the cisterns and the pial perivascular spaces.

**Fig. 2:**
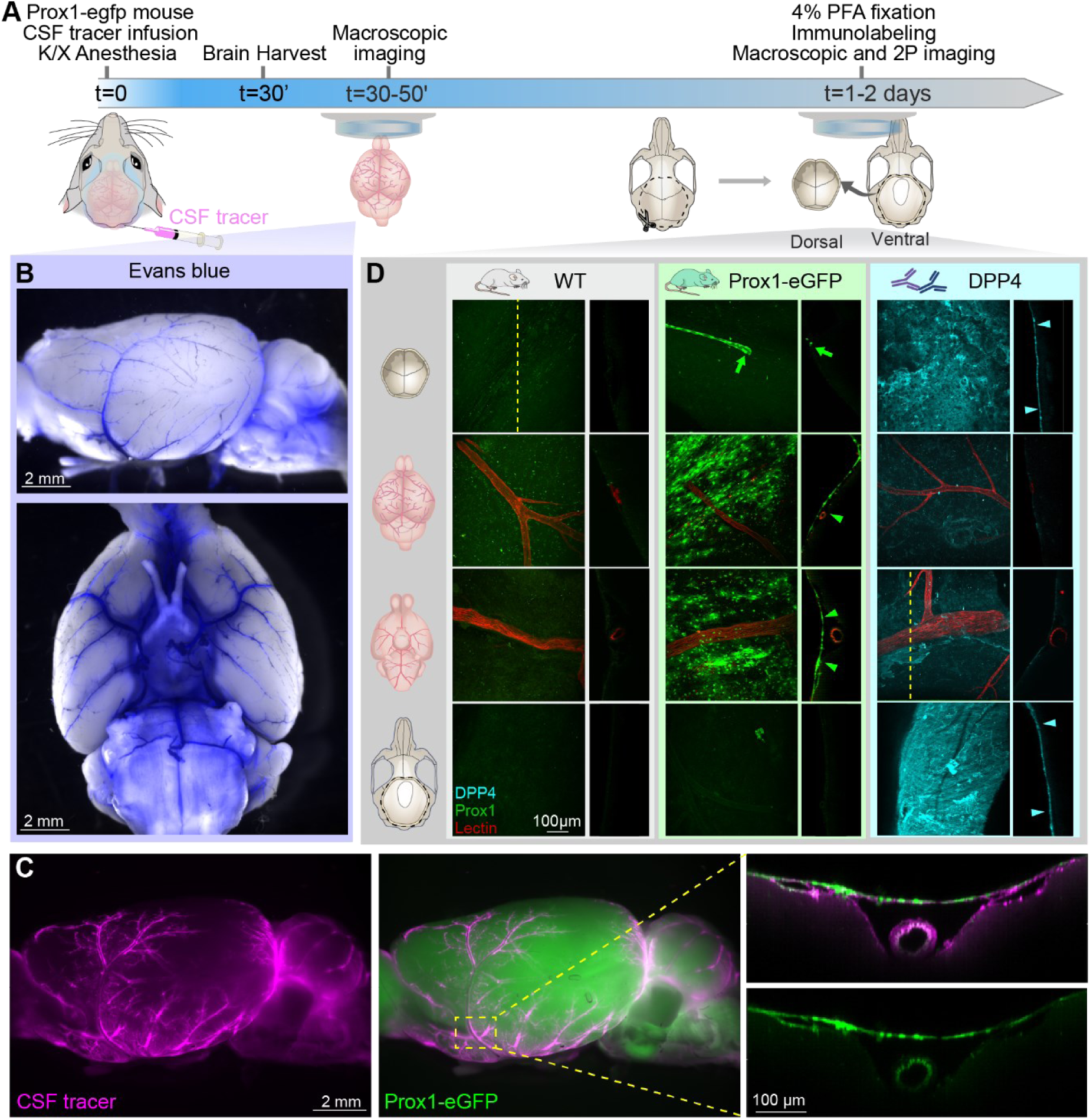
SLYM defines the subarachnoid cisternal system and the pial perivascular space. **(A)** Schematic of the experimental approach. CSF tracers, either Evans Blue (0.9 kDa, 0.5% in aCSF w/v) or BSA-647 (67 kDa, 0.5% w/v) were delivered (10 µL, 2 µL/min) through a cisterna magna cannula in K/X-anesthetized Prox1-eGFP mice. After 30 minutes of tracer circulation, the PROX1-eGFP mice were perfused with PBS to remove blood signal interference. The brains were imaged immediately after harvesting using a fluorescent macroscope. **(B)** Macroscopic fluorescent images of the whole brain showing the cisterns and pial perivascular spaces filled with a small size tracer, Evans blue (blue). **(C)** Representative images of a brain from an animal injected with a larger tracer, BSA-647 (magenta). The green signal corresponds to Prox1-eGFP expression. Close-up images over a major cerebral artery (middle cerebral artery) illustrate the accumulation of the fluorescent tracer below the Prox1-eGFP+ cells. **(D)** Skull and brain from both Prox1-eGFP and wildtype mice taken before and after DPP4 staining. Confocal and two-photon XY projections from whole-mount dorsal skull (first row), dorsal brain (second row), ventral brain (third row), and ventral skull (bottom row). Orthogonal projections were obtained as indicated by the dashed line in the first panel, except for the ventral brain after DPP4 staining where the projection was obtained from an area including a DPP4 positive section close to the vessel. A meningeal lymphatic vessel positive for Prox1-eGFP is observed in the dorsal skull of the Prox1-eGFP mouse (green arrow). The analysis showed that SLYM in the Prox1-eGFP reporter mice (green arrowheads) adheres to the brain surface, whereas the ABC layer, immunolabeled for DPP4 (cyan arrowheads), adheres to dura on the skull.

### High resolution immunohistochemical characterization of SLYM

The first description of SLYM (12) included a detailed immunohistochemical characterization of the newly discovered meningeal layer surrounding the mouse and human brain. To further characterize SLYM, whole heads along with the attached upper cervical region harvested from Prox1-eGFP mice were decalcified. Decalcification is the only approach that allows the preparation of thin paraffin sections, containing all the meningeal layers located between the parenchyma and the surrounding skull or vertebrae. Bright field immunohistochemistry based on horseradish peroxidase staining (HPS) was utilized rather than fluorescence labeling to ensure optimal sensitivity and specificity of the antibody labeling. Consecutive sections were used when comparing the pattern of labeling of multiple antibodies to avoid double labeling of the thin membranes (**Fig. 3-7**).

**Fig. 3:**
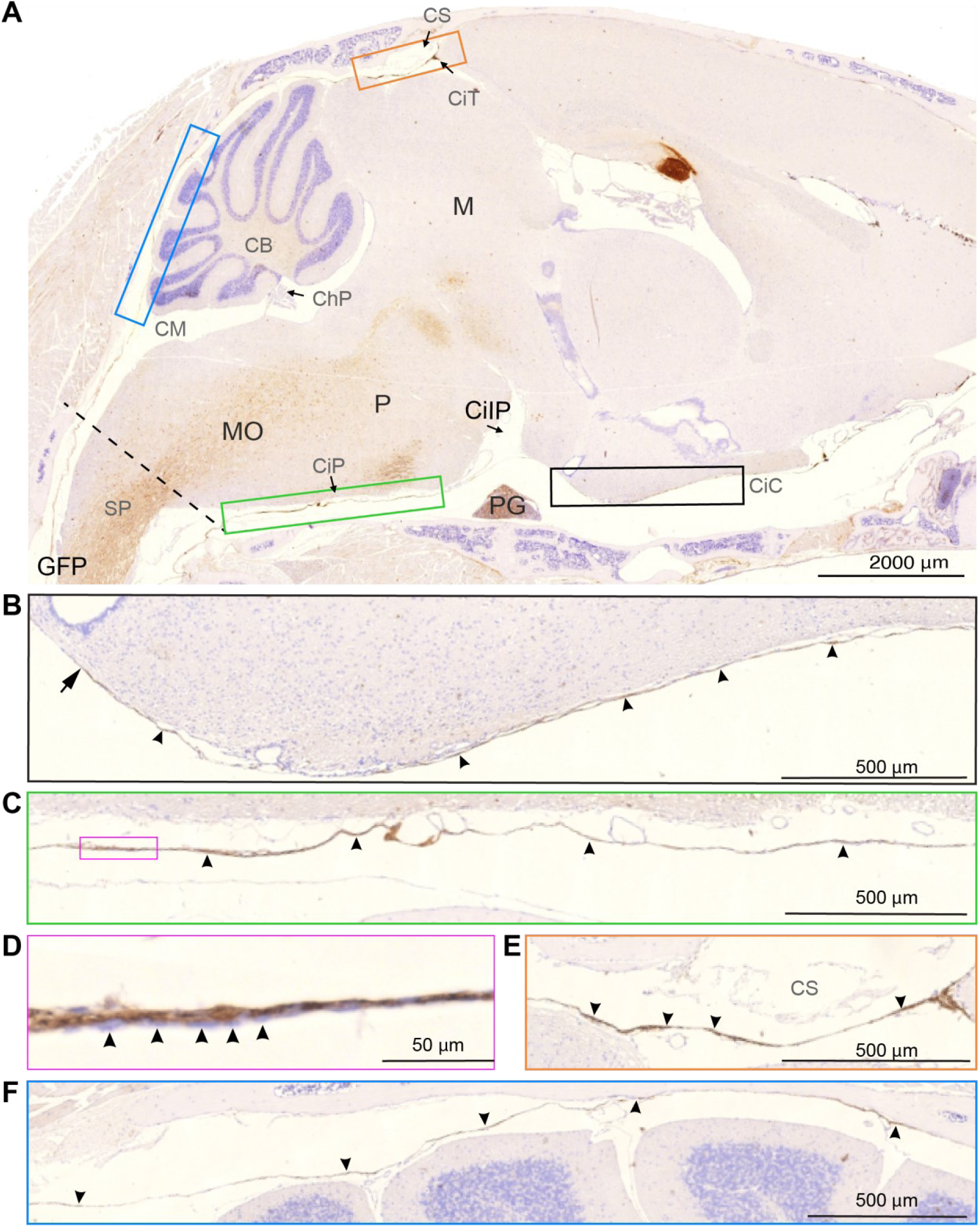
SLYM encases the brain parenchyma here shown in a sagittal section extending caudally from the upper spinal cord to the olfactory bulb region rostral. **(A)** GFP immunosignal in Prox1-eGFP mouse tissue forms a lamina covering the entire brain, delimiting the cisternal spaces. Colored insets are used to show high magnification images of different anatomical regions. The dashed line indicates the plane through foramen magnum separating the spinal cord (SP) from medulla oblongata (MO). **(B) Black insert** shows the rostral extension of the basal cistern (arrowheads). Note that the histochemical reactivity is attenuated corresponding to the median eminence (arrow). Cisterna interpeduncularis (CiIP) has been already analyzed in a previous publication (Møllgård, et al., 2023 (12), suppl fig.3). **(C) The green insert** depicts the cisterna pontis (CiP), lining the basal CSF space. **(D)** In a higher magnification (**magenta insert**) SLYM, with individual GFP positive cells (arrowheads), seems to be partly covered by an inner cell layer to the left. **(E) Orange insert** shows the region of confluence of sinuses (CS) where the GFP signal is clearly visible **(arrowheads),** delimiting the cisterna tegmentalis (CiT). **(F) Blue insert** shows the superior cerebellar cistern (cisterna vermis), delimiting two CSF spaces adjacent to the cerebellum. The one-layered delicate SLYM membrane deviates between attachment to the skull and the cerebellar pial surface. CB: cerebellum, ChP: choroid plexus, CiC: carotidand chiasmatic cistern, CiIP: cisterna interpeduncularis. CiP: cisterna pontis, CiI: cisterna tegmentalis, CM: cisterna magna, CS: confluence sinuum, M: mesencephalon, MO: medulla oblongata, P: pons, PG: pituitary gland, SP: spinal cord. Magnification is indicated by labelled bars on the individual figures.

**Fig. 4:**
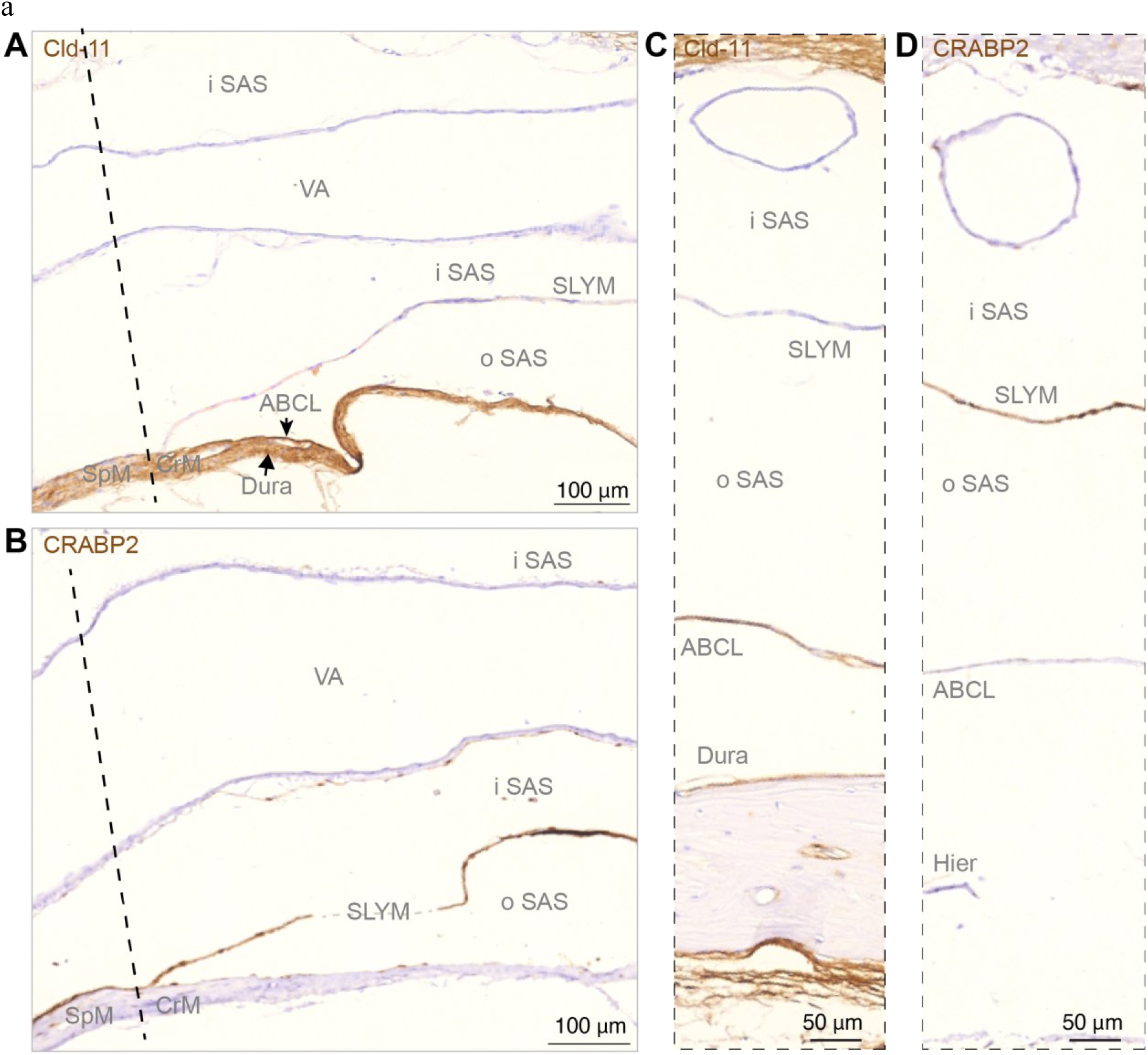
Sagittal sections show SLYM is not fused with the arachnoid barrier layer along basal cisterns. **(A, B)** Immunohistochemical analysis of arachnoid barrier cell layer (ABCL) stained for Cld-11 and SLYM stained for CRABP2 at the transition from spinal meninges (SpM) to cranial meninges (CeM) indicated by dashed line, corresponding to the plane of foramen magnum. SLYM and ABCL are fused corresponding to the rostral-most spinal meninges, but the deviation of the two layers is characteristic of the beginning of cranial meninges creating an inner subarachnoid space (iSAS) containing the vertebral artery (VA) and an outer subarachnoid space, between SLYM and the arachnoid barrier layer. In **(C, D)** corresponding to cisterna pontis the arachnoid barrier cell layer (ABCL), positively reacting for Cld-11, covers dura but renders SLYM negative. SLYM shows positive immunoreactivity for CRABP2 whereas the ABCL is not stained. Due to the heat-induced epitope retrieval (Hier) a fraction of bone with dura disappeared from this section. Dashed line indicates a nearly frontal plane through foramen magnum separating the spinal cord from medulla oblongata and thus the cranial meninges (CrM) from spinal meninges (SpM). ABCL: arachnoid barrier cell layer, CrM: cranial meninges, Hiar: heat-induced antigen retrieval, iSAS: inner subarachnoid space, oSAS: outer subarachnoid space, SpM: spinal meninges, VA: vertebral artery. Magnification is indicated by labelled bars on the individual figures.

**Fig. 5:**
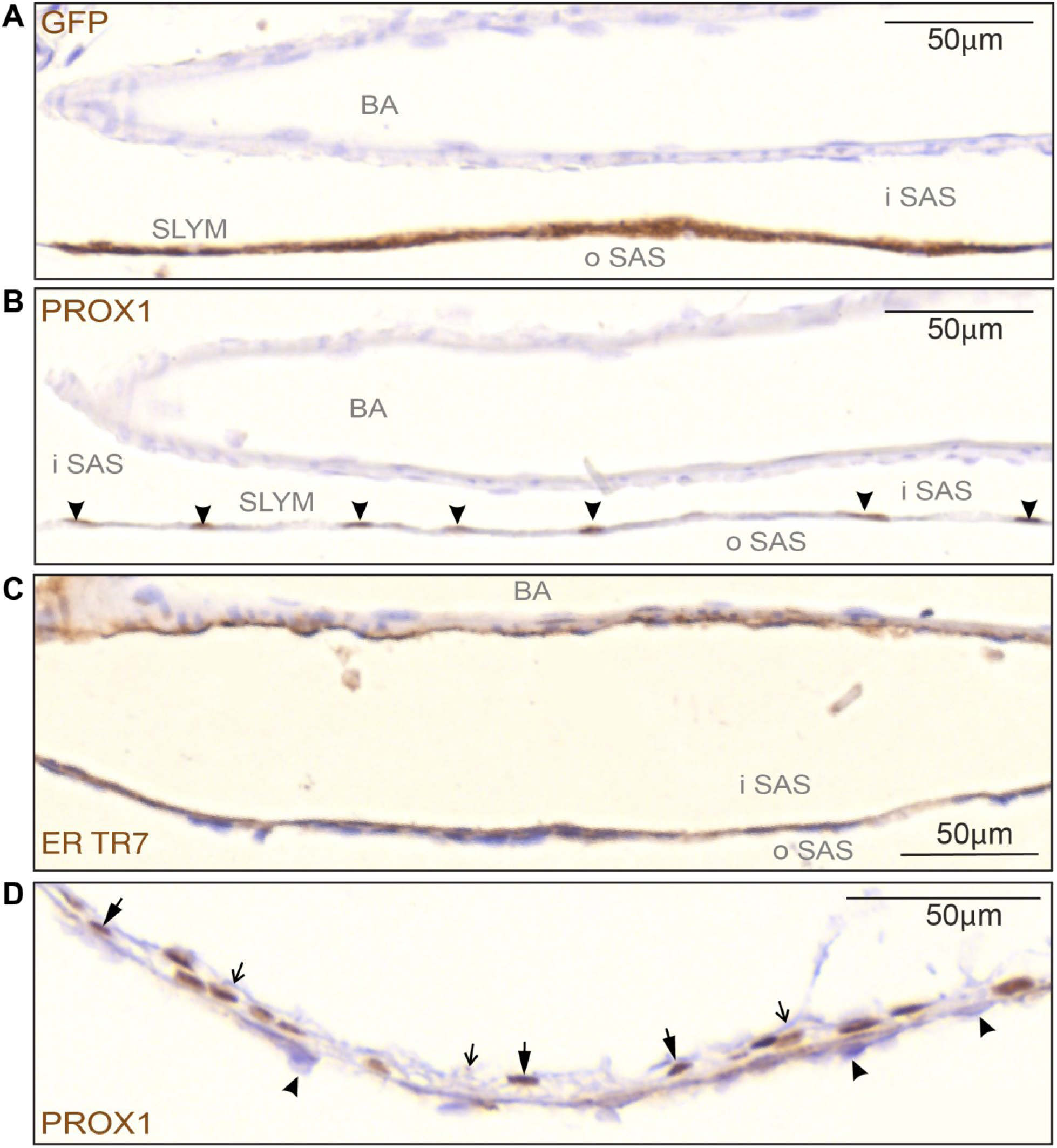
SLYM is characterized by a specific subset of immunological markers. **(A-C)** High magnification images of the basilar artery on the ventral part of the brain show the division of the subarachnoid space (SAS) into an outer and inner compartment by the SLYM meningeal layer (GFP+ labelled). **(B)** Prox1 immunolabeling of serial sections of the same tissue allowed individual identification of SLYM cells, due to their immunoreactive nuclei (arrowheads). **(C)** ER-TR7 immunoreactivity is found on the inner surface of SLYM, showing the presence of Collagen type VI on this meningeal layer. Also note the staining of adventitia of the basilar artery (BA). **(D)** The Prox1-stained nuclei (thick arrows) in the wall of cisterna ambiens are in direct contact with arachnoid barrier cells towards the exterior (arrowheads) and with an unstained layer of arachnoid reticular cells facing the inner subarachnoid space (thin, open arrows). BA: basilar artery, iSAS: inner subarachnoid space, oSAS: outer subarachnoid space. Magnification is indicated by labelled bars on the individual figures.

**Fig. 6:**
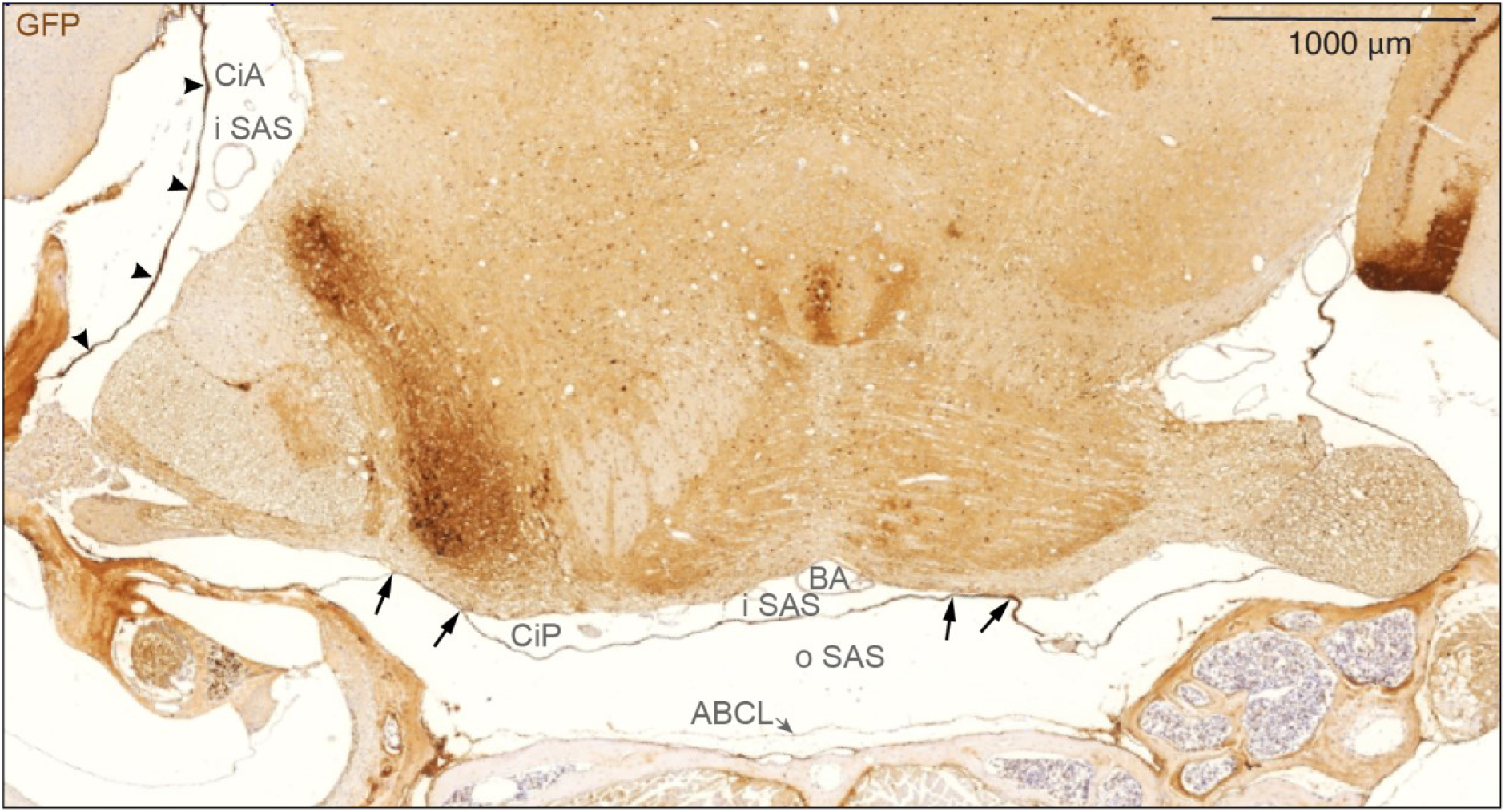
SLYM encases brain parenchyma evident in a coronal section through mid-pontine region. Immunohistochemical analysis of GFP signal of a coronal section of Prox1-eGFP tissue allowed additional characterization of the meningeal layers covering the parenchyma. SLYM (GFP+) layer divides the subarachnoid space into two well-defined compartments, an outer subarachnoid space (oSAS) facing the arachnoid barrier cell layer (ABCL) and an inner subarachnoid space (iSAS), enclosing blood vessels, here the basilar artery (BA). SLYM seems to fuse with pia covering a fraction of the basolateral part of pons on both sides (arrows) thus creating a sub-compartmentalization of the inner SAS. The strongly stained cisterna ambiens (CiA) is indicated by arrowheads. ABCL: arachnoid barrier cell layer, BA: basilar artery, CiA: cisterna ambiens, CiP: cisterna pontis, iSAS: inner subarachnoid space, oSAS: outer subarachnoid space. Magnification is indicated by labelled bar.

**Fig. 7:**
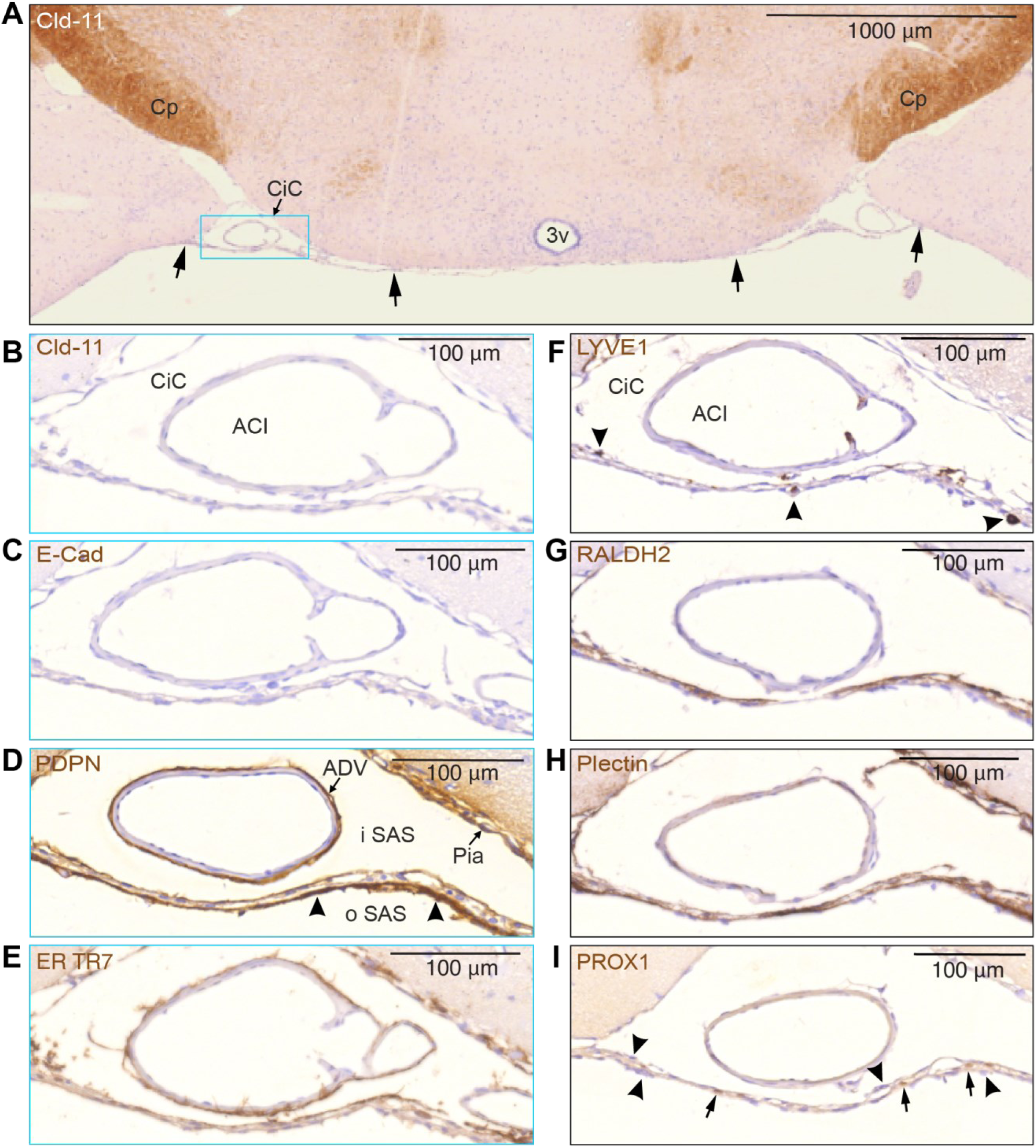
SLYM and the ABC layer can be differentiated by their distinctive immunomarker expression. **(A)** The unstained SLYM in this claudin-11 stained section located at a middle coronal position of the circle of Willis, at the suprachiasmatic level, indicated by arrows, seems to fuse with the pia along both medial and lateral borders of the cisterna caroticus (CiC) - blue rectangle - which is shown in higher magnification in **(B-I)**. Note the positive Cld-11 reactivity in the oligodendrocytes of the cerebral peduncle (Cp) in contrast to the negative appearance of SLYM in **(B). (C)** E-cadherin staining also provided a negative response in SLYM, pia and adventitia of the internal carotid artery (ACI). **(D-I)** show SLYM covered by an inner and outer layer of different types of arachnoid leptomeningeal fibroblasts. The PDPN-stained section **(D)** demonstrates a strongly reactive layer (arrowheads) facing the outer SAS (oSAS), a weaklier stained SLYM in the middle and yet another thin leptomeningeal layer facing the inner SAS (iSAS) with a positively reacting pia. The adventitia (ADV) surrounding the internal carotid artery (ACI in B) is also positively stained. **(E)** ER-TR7 immunoreactivity demonstrates collagen type VI in particular on the inside of SLYM but is also associated with adventitia of ACI and pia. **(F)** As expected by its “lymphatic-like” nature, SLYM acts as an immunological niche. LYVE1 antibody highlights perivascular macrophages (arrowheads) located either on the inner, middle or outer part of SLYM. **(G)** Retinaldehyde dehydrogenase (RALDH2) is particularly strongly stained along the inner SLYM layer. **(H)** Plectin, a possible key cytoskeleton interlinking molecule, is also present in SLYM meningeal lamina, in particular facing the outer subarachnoid space, but also staining the pia. **(I)** PROX1-stains nuclei (arrows) identify SLYM as the middle layer in this triple-layered structure where nuclei in inner and outer layers are PROX1-negative (arrowheads). 3v: third ventricle, ACI: internal carotid artery, ADV: adventitia, CiC: cisterna caroticus, Cp: cerebral peduncle. Magnification is indicated by labelled bars on the individual figures.

eGFP immunolabelling was first employed to identify the exact anatomical location of SLYM in sagittal sections in PROX1-eGFP reporter mice (**Fig. 3**). The analysis showed that SLYM forms a meningeal layer encasing the entire brain, including the ventral and dorsal brain surfaces (**Fig. 3A**). Higher power micrographs of the sagittal section document that SLYM forms a roof covering the rostral basal cisterns (**Fig. 3B**), the more caudal cisterna pontis (**Fig. 3C**), and dorsally delineates cisterna tegmentalis (also named cisterna superior), as indicated by arrowheads (**Fig. 3E**), and the superior cerebellar cistern (**Fig. 3F**). Note that the slender and more delicate SLYM membrane covering the superior surface of cerebellum deviates from attachment to the skull (two right arrowheads pointing upwards) to approach the cerebellar pial surface (three left arrowheads pointing downwards). The somewhat thicker appearing SLYM of the cisterna pontis (**Fig. 3C**), part of which is shown in higher magnification in **Fig. 3D**, consists of two layers – the one-layered SLYM *per se* adjoined by an outer unstained cell layer (arrowheads).

We next ask whether Prox1-eGFP+ SLYM cells in certain regions merged to form a double membrane with the ABC layer. For this analysis, the well-characterized marker for the ABC layer, an antibody directed against the tight junction protein Claudin-11 (24), was used in combination with CRABP2 that labels SLYM (12, 15). Upon examining the relationship between the two membranes, it became evident that SLYM and ABC layers are fused below foramen magnum, and together form a double membrane adjoining dura, i.e., spinal meninges that covers the cavity of the upper spinal canal (**Fig. 4A**). However, rostral to the foramen magnum, corresponding to cranial meninges, SLYM and ABC layers separate: the ABC layer is located in close proximity to dura, while SLYM continues as solitary membrane subdividing the subarachnoid space (SAS) into an inner and outer CSF-filled compartment at the base of the brain (**Fig. 4B**). Inner and outer subarachnoid spaces are obvious at the mid-pontine level (**Fig. 4C-D**).

High magnification imaging of SLYM around the basilar artery (**Fig. 5A-C**) provided additional evidence for compartmentalization of the SAS by SLYM. Serial sections prepared from a Prox1-eGFP reporter mouse were reacted with multiple antibodies, including eGFP, Prox1 (note the nuclear labeling – arrowheads in **Fig. 5B**), and ER TR7, which also labeled the adventitia of the artery (**Fig. 5C**). SLYM from cisterna ambiens (**Fig. 5D**) is closely attached to the ABC layer (arrowheads), but is also coherent with an unstained layer of arachnoid reticular cells facing the inner subarachnoid space (thin, open arrows).

Immunostaining of coronal sections against eGFP prepared from a Prox1-eGFP reporter mouse depicted again that SLYM forms a complete membrane that covers the basal cisterns (here cisterna pontis (CiP)) and is separated from the ABC layer (**Fig. 6**). SLYM seems to fuse with pia along the basolateral pons, thus creating separated inner SAS spaces – one median and two lateral (see arrows in **Fig. 6**). The wall of the lateral cisterna ambiens (CiA) is always thick and strongly stained.

Additional immunostaining showed that SLYM also fuses with pia along the medial and lateral borders of the cisterna caroticus (CiC) (arrows in **Fig. 7A**). This figure which is stained for claudin-11 demonstrates that SLYM is distinct from and not stained by anti-claudin 11. **Fig. 7B-I** depict SLYM covered by inner and outer layers of fibroblasts characterized by several different antigens. The detailed analysis of serial sections documented that SLYM *per se* is negative for claudin-11 and does not stain positive for E-Cadherin or LYVE1. However, SLYM and associated fibroblast layers exhibited positive immunolabeling for PDPN, ER TR7, RALDH2, plectin and Prox1. Additionally, the analysis showed that SLYM hosts several LYVE-1 immunopositive macrophages (**Fig. 7F**).

In our fixed histology material, we have encountered the one-layered SLYM membrane as an entity by itself (**Fig. 4**), as well as SLYM covered by or covering another layer of leptomeningeal fibroblasts, thus appearing as a part of a two-layered structure (**Fig. 3D**) as our previously demonstrated ABC/SLYM layers. In certain sub-regions of the basal cisterns, SLYM appears to be ‘sandwiched’ between both an inner and an outer lepto–meningeal fibroblast layer (**Fig. 7**). The presence of ‘sandwiched SLYM’ in the basal cisterns raises the possibility that extra leptomeningeal layers are required to support and reinforce SLYM in this region due to the heavy pulsations from the large basilar arteries.

### Comparing new data with the classical literature

Has a meningeal membrane matching SLYM been described in the classical literature? To interrogate previously published data, we reproduced the toluidine blue-stained semithin section-based scheme of Orlin et al. in 1991 (21). In fact, the Orlin publication reproduced the classical sketch from Nabeshima et al. in 1975 to frame their own data into a historical context (20). We have here color-coded the panels from both publications to compare the meningeal layers in these publications to the data reported here (**Fig. 8A-B**). The Nabeshima scheme, as well as the Orlin semithin section, depicts a mono-cell layer (green) between the arachnoid and the pial cell layers (both gray) that covers a pia-adjacent vessel (red). We propose that the green-colored membranes in **Fig. 8A-B** represent SLYM, as these membranes rest on the pia-adjacent vasculature and the cytosol of the cells is translucent. The panels depict the green membrane as separating the SAS into two compartments and clearly illustrate that this subdivision of SAS differs from the artificial subdural space between dura and the ABCL in the Nabeshima scheme. For comparison, an orthogonal view of a pia-adjacent blood vessel containing CSF tracer in the perivascular space from our analysis is included (**Fig. 8C**). A graphical scheme is included on top to illustrate that the SLYM layer defines the pial perivascular space by creating a roof resting on top of the pia-adjacent vasculature (**Fig. 8C**).

**Fig. 8:**
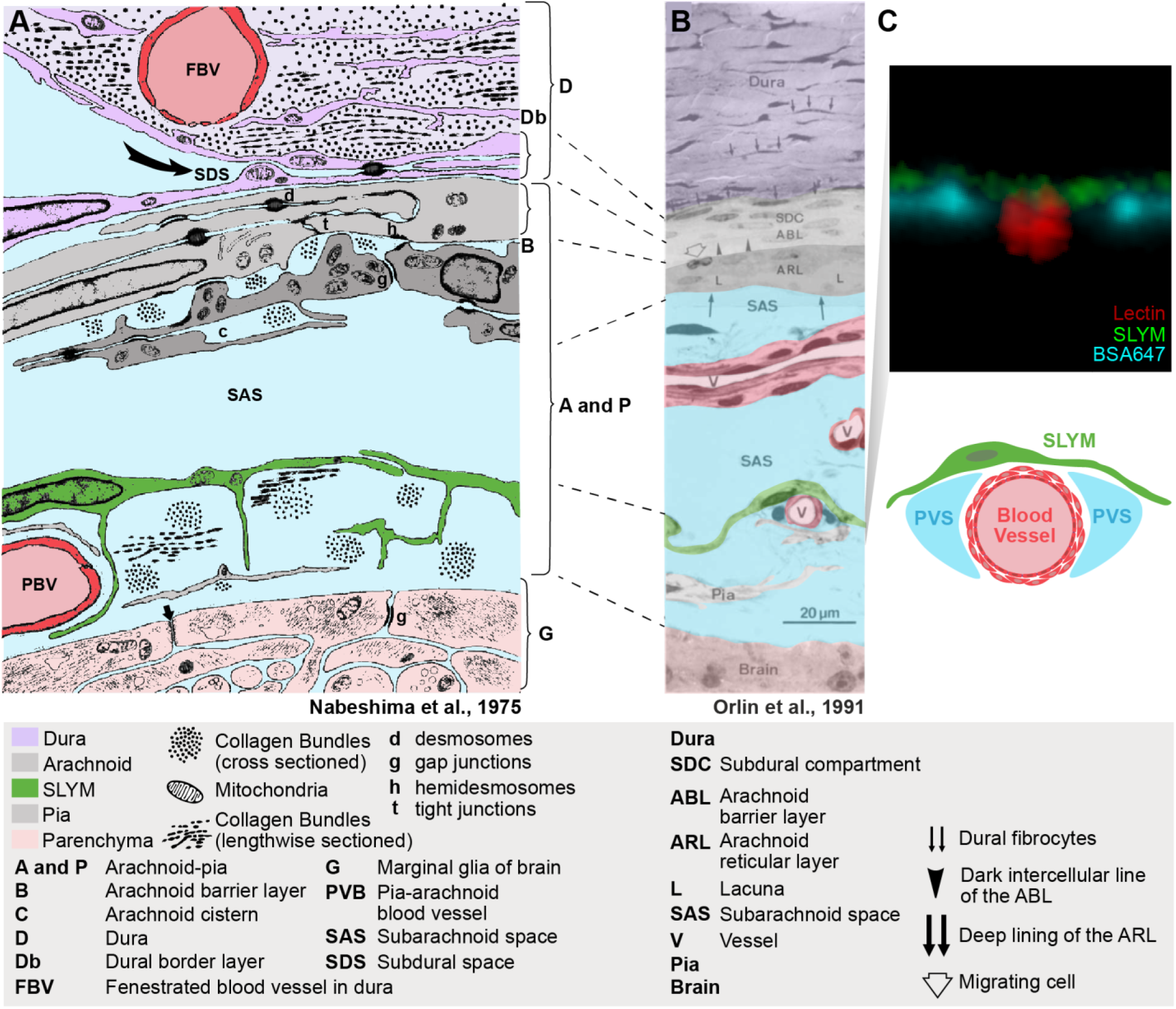
SLYM is illustrated as a distinctive meningeal layer in the classical literature. **(A)** Reproduced sketch from Nabeshima et al., 1975 (20). The sketch is based on examination of the meningeal region in studies of multiple mammals (mice, rats, chinchillas, rabbits, cats and Macaque monkeys) by electron microscopy and freeze-fracturing. Meningeal layers have been color-coded for clarity, without changing the original sketch. SLYM layer is already indicated as an intermedial layer between the arachnoid (ABC and ARLlayers) and pia, resting on top of a pial vessel (PBV). **(B)** Light microscopy of a thin section stained with toluidine of meninges in the Norwegian landrace pig image from Orlin et al., 1991 (21). Orlin et al., compared this section to the sketch in the Nabeshima publication and the original labels and lines have not been changed. **(C)** Confocal cross-section of a pial vessel from this study (Fig. S2E) is included to compare the structural details with our observations. The sketch below illustrates that SLYM forms the roof of the pial perivascular spaces.

## Discussion

The meninges surrounding the brain are classically identified as dura, arachnoid and pia. The meningeal layers are all composed of connective tissue and their designation is based on macroscopic postmortem examination. The simplified classification of the meningeal membranes into 3 layers was questioned already in the first detailed histological studies of the meninges (20, 21). We recently described a 4^th^ meningeal membrane, SLYM (<u>S</u>ub-arachnoid <u>Ly</u>mphatic-like <u>M</u>embrane), in both mouse and human brains (12). SLYM is located within the subarachnoid space (SAS), the large CSF-filled spaces that surround the brain, between the arachnoid and pia (12). We here reexamined the structural characteristics of SLYM. We confirmed that SLYM surrounds the entire brain and, based on immunolabeling, demonstrated that SLYM phenotypically differs from the arachnoid barrier cell (ABC) and pia layers (**Figs. 2-6**). Our findings revealed that SLYM rests on top of the pia-adjacent vasculature, forming a roof over the cisterns and the pia-adjacent vasculature. As such, SLYM creates a continuous CSF-filled space encompassing the cisterns and the pial perivascular spaces. The unique position of SLYM becomes particularly evident after extracting the brain in Prox1-eGFP reporter mice wherein a CSF tracer circulated for 30 min. Macroscopic imaging of freshly resected brains showed that SLYM retains the CSF tracers within the cisterns and the pial perivascular spaces, outlining a large tracer-filled space that limits the spread of both a small and a large CSF tracer. When dissecting the brains of Prox1-eGFP reporter mice, we observed that SLYM preferentially adheres to the brain’s surface rather than to the dura, which remains firmly attached to the skull. Conversely, the ABC layer, marked by positive DPP4 immunolabeling, exhibits a distinct adherence to the dura and skull. Similar separation between SLYM and the ABC layer is also evident in decalcified, paraffin-embedded sections (**Fig. 1**). Hence, these observations confirm that SLYM divides the larger subarachnoid space into two separate compartments (**Fig. 2**).

The subarachnoid space (SAS) is in most textbooks depicted as a single compartment. However, Liliequist (25) described already in 1956 a compartmentalization of the SAS. He described a ‘*thin arachnoid membrane*’, considering this layer as ‘*the line of demarcation between the chiasmatic and the intercrural cisterns, possessing as it does the advantages of a definite anatomic structure*’. This membrane, which has become an essential intraoperative landmark named after its discoverer (26), is one of the better-known subdivisions of the SAS, but not the only one (27).

In the past, special attention has been given to the ABC layer, often described as a single barrier membrane that isolates the brain from the entrance of exogenous substances. Brøchner et al., 2015 (24) identified claudin 11 as the tight junction protein responsible for the barrier function of the ABC layer. However, multiple studies have documented that the ABC layer is more complicated, consisting of multiple cell types based on their ultrastructural features. In 1974, Lopes and Mair (28) summarized in their publication an analysis of the meningeal membrane in diverse animals and concluded that the arachnoid membrane is formed by at least two layers of cells, the outermost of them being electron dense with tight junctions and accompanied by an inner layer of clear cells occasionally separated by spaces and surrounded by loose collagen fibers. A basement membrane was noted to separate the superficial dark cells from the inner layer of the arachnoid membrane. In addition, dense cells with features of fibroblast could be found along the collagen fibers of the inner layer of clear cells (28). Nabeshima et al. 1975 conducted an electron microscopy (EM) and freeze-fracturing study, which was also based on analysis of multiple species. This publication also concluded that the ABC layer differs from the inner arachnoid cells with lighter cytosol (20). Moreover, Nabeshima et al, stated that: “*The cells of the pia were similar to the arachnoid cells but their processes were somewhat thinner. We were unable to find any cytological criteria for differentiating these cells from the layer of arachnoid cells next to the pia and, therefore, consider this inner layer of arachnoid and the pia to be basically one layer in which the different shapes of the cells are determined by the shapes of the structures they invest”.* This note highlights the fact that the classification of the 3 connective tissue meningeal membranes, dura, arachnoid and pia is based on macroscopic examination and that the inner arachnoid and the outer pial layers in histological sections do not exhibit distinct characteristics. The Nabeshima publication included a schematic of the meningeal layers, in which only features common to all the specimens of animals studied were included (20). This schematic was later used by Orlin et al. in 1991 (21) to illustrate the similarities between their own findings to those of Nabeshima et al. (20). We have here reproduced the figure from the Orlin et al publication along with data from our own study to highlight the similarities between the 3 reports (**Fig. 8**).

In 1983 - 84, Krisch, Leonhardt and Oksche’s work (29, 30) moved the field forward by studying the functional role of this meningeal segmentation through the injection of horseradish peroxidase (HRP) in different regions of the brain and meninges. They described that HRP distributed within at least 2 separate compartments of the SAS, which they entitled the arachnoid or pial spaces. The authors also defined the ‘*intermediate lamella*’, a structure composed by the inner arachnoid layer and the outer pial membrane, that dissociates when joining a blood vessel bridging in between the dura to the pia-adjacent vasculature. The intermediate lamella here forms a perivascular space surrounding the bridging vessel (30). HRP injected into either the arachnoid or pial spaces demonstrated that the intermediate lamella is indeed a barrier that separates the CSF-filled arachnoid and pial compartments, as we later described (12). The same technique was used by Orlin et al. 1991 (21), who infused HRP-mixed blood to mimic acute subdural hemorrhages. Their detailed electron microscopy analysis led to the description of the arachnoid reticular cells, as ‘*a 10-20 µm thick layer composed of irregularly branched cells’*. Adhering junctions were noted scattered both between the arachnoid reticular light cells and between these and the barrier cells (21). Later, Vandenabeele, Creemers and Lambrichts 1996 (31) described the ‘*arachnoid border cells*’ as sinuous flattened cells found along the inner surface of the arachnoid reticular layer, with uncertain nature, that could belong to either the arachnoid or pial layer. Several investigators have noted that the cytoarchitecture of these cells is so similar that the cells cannot be designated as belonging to either the inner arachnoid or outer pia cell layers based on their cellular characteristics.

Recently, Pietilä et al., 2023 confirmed the presence of a Prox1+ cell layer below the ABC layer (19). Electron microscopy and transcriptome analysis added additional support to the presence of a distinct Prox1+ cell layer which cells were coupled by adherence junctions composed of ve-Cadherin (CDH5) and gap junctions, Cx43, Cx30 and Cx26 (Gja1, Gjb6 and Gjb2). The authors documented that a Prox1-tdTomato+ layer is located below the ABC layer. A publication by the same groups interrogated the location of SLYM in relationship to the ABC layer and pia (18). *In vivo* analysis showed that SLYM (Prox1-tdTomato+/ve-Cadherin-GFP) is separated from dura by a CSF-filled space in cortex (Figs. 1L and 6F). However, the authors concluded that the CSF-filled space was artificial and created during surgery or injections. This conclusion was based on the evidence that depicts the meningeal layers adhere to each other in histological sections, and the observation that the depth of CSF-filled space appears to enlarge during imaging and depends upon the surgery (18). One problem with the *in vivo* experiments was the use of ve-Cadherin reporter mice that are not specific to any of the meningeal layers. In the ve-Cadherin-eGFP reporter mice, a subpopulation of cells located in both dura and the ABC layer express GFP, as well as most cells in SLYM and pia. Thus, the GFP signal functions as a pan-meningeal label and it is not possible to separate the distinct meningeal layers *in vivo*. This conclusion is supported by the single cell transcriptome analysis showing that dural border cells, ABC layer, SLYM (Prox1-positive cells named BFB2 and 3) and pial cells all express ve-Cadherin (19).

We have in this study used markers that are only expressed by either the ABC layer or by SLYM. That enabled us to demonstrate, both in the freshly resected brain (**Fig. 2**) and *ex vivo* (**Figs. 3-7**), that the subarachnoid space is subdivided into two separate compartments by SLYM. SLYM acts as a barrier for a CSF small tracer (Evans blue, 1 kDa), a slightly larger tracer (TMR-conjugated dextran, 3 kDa), a large tracer (or BSA-647, 67 kDa) and 1 µm microspheres (**Fig. 2**) (12). It is here important to note that the literature is based on postmortem histological analysis, which results in major displacement of the CNS structures (11). The MR imaging of the live brain *in situ* when compared with paraffin sections of whole head provided direct evidence for major restructuring of the SAS due to the shrinkage of the soft tissue (brain, meningeal membranes, vasculature and more), while the structural properties of bony skull do not notably change (**Fig. 1**). We propose that the fragile SLYM layer located in the SAS is displaced during the specimen preparation. In histological sections, SLYM seems to adhere to either the ABC layer or pia, depending on where the fixative first reaches the tissue and induces shrinkage of the subarachnoid space and deformation of the leptomeningeal membranes.

In general, SLYM appears as a single, thin cell layer that often seems to fuse with the E-Cadherin positive ABC layer membrane over dorsal cerebral cortex in histological sections, as documented in Møllgård et al. (Fig. 4C-D in ref (12)) and originally described by Orlin et al. in the spinal cord (21). A similar appearance is present at the cerebellar surface. In the caudal cisternal system, SLYM is a part of a two-layered membrane covered towards the inside by a collagen-VI containing fibroblast layer. Around the rostral basal cisterns, however, SLYM is ‘sandwiched’ between an inner and an outer fibroblast layer creating a thick 3-layered membrane clearly separated from the ABC layer (**Fig. 7**). RALDH2 exemplifies the inner layer and plectin the outer. The reinforcement/strengthening of SLYM covering the cisterns containing the large intracranial arteries might reflect that arterial pulsatility in these cisterns drives fresh CSF from the ventricles up along the anterior, middle, and posterior cerebral arteries. At the transition between the skull and the vertebrae, SLYM appears to fuse with the ABC layer (**Fig. 3**)

As discussed, the best separation of SLYM and the ABC layer was noted at the base of the brain. SLYM here serves the important function of separating the subarachnoid space into two CSF-filled compartments facilitating the transport of newly produced CSF along the perivascular spaces. Loss of the SLYM barrier function here may be expected to disrupt the unidirectional glymphatic transport of CSF, with the result that CSF containing amyloid, tau, synuclein and other waste products is recirculated back into the brain. The resultant inflow of aggregation-prone peptides and proteins may contribute to both the initiation and progression of proteinopathic neurodegenerative diseases. Future studies dissecting key questions, such as the age-related atrophy of SLYM, or its role in neuroinflammation, will be necessarily imprecise in the absence of a designated name for the SLYM membrane. The existing names for SLYM include the inner arachnoid layer, the reticular layer of the arachnoid membrane, and latest, BFB2-3. We propose that the name SLYM, used to specify a distinct meningeal membrane, will help to ensure the precision and replicability of future studies – in an unnecessarily contentious field.

## Materials and Methods

### Animals

Both wildtype C57BL/6JRj and Prox1-EGFP+ (Janvier Labs, Le Genest-Saint-Isle, France) (14) female and male mice were utilized at postnatal age 90-120 days. Mice were housed in groups, containing 4-5 mice per cage, with controlled humidity and temperature, 12 hours light-dark cycle (6:00 AM/6:00 PM) and regular access to tap water and chow food ad libitum. All experiments carried out at the University of Copenhagen were approved by the Animal Experiments Council under the Danish Ministry of Environment and Food (license number: 2015-15-0201-00535) and the procedures were performed in accordance with the European directive 2010/63/EU, with every effort undertaken to reduce the utilization of animals. Experiments conducted at the University of Rochester Medical Center were approved by the University of Rochester Committee on Animal Resources.

### MRI scans

Scans were performed on a 9.4 T animal scanner (BioSpec 94/30USR, Bruker BioSpin, Ettlingen, Germany) equipped with a cryogenically cooled quadrature-resonator (CryoProbe, Bruker BioSpin). Head movement during scanning was minimized by restraining the animals on an MR-compatible stereotactic holder with ear bars. During MRI scanning, the animal was anesthetized under 1-1.5% isoflurane in a 1/1 mixture of air/oxygen delivered via nose cone. Cardiac and breathing rate was monitored, and body temperature was maintained at 37 °C for the duration of the experiment of live imaging (SA Instruments). For brain and CSF volume measurement, a 3D constructive interference in steady-state (3D-CISS) image was calculated as a maximum intensity projection (MIP) from 4 realigned 3D true fast imaging with steady precession (3D-TrueFISP) volumes with 4 orthogonal phase encoding directions (repetition time = 4 ms, echo time = 2 ms, repetitions = 2, flip angle = 40°, spatial resolution 75 µm × 75 µm × 75 µm, RF phase advance 0, 180, 90, 270°, total acquisition time = 46 minutes) (32). After the live imaging session, the animal was allowed to inspire 100% nitrogen delivered via a nose cone, which was firmly placed around the mouse’s snout, and cardiorespiratory arrest was confirmed (22). The 3D-CISS scanning was repeated at 1, 2, 6, 12, 18, and 24 h after cardiorespiratory arrest. To obtain optimal spatial uniformity, all acquired 3D-TrueFISP volumes were motion-corrected before calculation as MIP, and the image bias field was removed with the N4 bias field correction algorithm of SimpleITK (33, 34). For each brain sample, the total brain volume was automatically segmented by using region growing with ITK-snap (version 3.8.0) (35). In addition, the pixel intensity factorized semi-automatic thresholding was performed to segment the CSF space (including ventricles and subarachnoid spaces) in each hemisphere. The segmentation and volume measurements were performed in Imalytics Preclinical software (ver. 3.0.2.5, Gremse-IT GmbH, Aachen, Germany).

### Cisterna Magna Infusion

Mice were anesthetized by injecting ketamine and xylazine mixture (K/X: 100 mg/kg, 20 mg/kg) intraperitoneally (i.p.). Additional anesthesia (K/X: 50 mg/kg, 10 mg/kg) was administered as needed, accompanied by ongoing maintenance and monitoring body temperature at 37 °C. The anesthetized Prox1-eGFP mice were placed in a stereotaxic frame, with the head tilted down in concord position. Then, the skin covering the back of the neck was shaved and opened to visualize the underlying muscles, and dissected to visualize the cisterna magna. A 30G-needle attached to a PE-10 line containing either Evans Blue [0.9kDa, 0.5% (w/v) in aCSF] or Bovine Serum Albumin tagged with Alexa 647 [67kDa, 0.5% (w/v) in 2% gelatin in aCSF] was then inserted and secured in place. After detaching the animals from the stereotaxic frame and allowing the recovery of the natural laying down posture, 10 µL, 2 µL/min, were infused into the cisterna magna with a syringe pump (Harvard Apparatus) as described before (36). 30 minutes after the start of the infusion, animals were transcardially perfused with 20 mL of PBS to eliminate any blood present in the tissue and brains were quickly extracted. Tracer influx into the brain was imaged ex vivo by macroscopic whole-brain (Leica M205 FA with a Hamamatsu ORCA-Flash4.0 V2 Digital CMOS camera using LAS X Leica software).

### Whole mount brain and skull immunohistochemistry and imaging

Both wildtype C57BL/6JRj and Prox1-EGFP+ mice were anesthetized using ketamine/xylazine (100/20 mg/kg) and the cisterna magna was cannulated for tracer infusion (10µL, 2µL/min, 70kDa Texas Red-Dextran. 0.5% (w/v) in aCSF). The tracer was allowed to circulate for 30 minutes, and 5 minutes before the sacrifice, 100mL of Wheat Germ Agglutinin conjugated to AlexaFluor-647 (1mg/mL in PBS) was injected retro-orbitally to label the vasculature. Mice were then perfused with 20mL ice cold PBS, then brains were carefully dissected to preserve the dura membrane on the skull surface, and SLYM membrane on the brain surface. Brains, skulls, and livers were immersion fixed in 4% paraformaldehyde in PBS overnight at 4C.

For immunohistochemistry, whole brains were cut in half sagittally, and muscle and skin dissected off skulls. Whole mount brains and skulls underwent the same immunohistochemistry treatment. Samples were washed 3 times in PBS and blocked at room temperature for 1hr in blocking solution (5% Normal Donkey Serum, 0.3% Triton, 3% Fab, 0.2% Gelatin in PBS). The samples were then washed 3 times in PBS, then immersed in primary antibody solution overnight on an orbital shaker at 4C (5% Normal Donkey Serum, 0.3% Triton, 0.2% Gelatin, 1:200 goat anti-Dpp4 (R&D Biosystems, AF954)). Samples were washed 3 times in PBS and incubated in secondary antibody solution for 2 hours at room temperature on an orbital shaker (5% Normal Donkey Serum, 0.3% Triton, 0.2%Gelatin, 1:500 donkey-ani-goat-Cy2 (705-225-147 Jackson Immunoresearch)). Samples were then washed 3 times in PBS and imaged immediately.

Brains and skulls from wildtype mice stained for DDP4 were first imaged on a fluorescent macroscope (LAS-X software; Leica, M205FA). Then brains and skulls were placed in a 2 well chamber with glass coverslip bottom (thickness #1.5H, 80287, Ibidi) and filled with PBS solution. Confocal images were acquired with an Olympus Fluoview microscope (Objective: 10x 0.4N.A. Olympus), and Z-stacks were acquired of dorsal and ventral brain and skull surfaces. 2-Photon images were acquired with a resonant scanner B scope (Thorlabs), a 20x Objective (1.0N.A. water dipping, Olympus) and a tunable pulsed laser (Chameleon Ultra II, Coherent). Sequential z-stacks were acquired (ThorImage) at 890nm for second harmonic excitation of collagen fibers, and at 820nm for simultaneous excitation of WGA-647, TxRed-Dextran, and Cy2 antibody. Sequential stacks were aligned using ImageJ software. Prox1-eGFP positive mice were imaged identically, but not stained for DPP4.

### Immunohistochemistry of paraffin-embedded sections

Paraffin-embedded sections from C57BL/6JRj mice were processed according to standard protocols. In brief, endogenous peroxidase activity was quenched and non-specific binding was inhibited by incubation for 30 minutes with 10% goat serum (Biological Industries, Kibbutz Beit-Haemek, Israel) at RT. Sections were incubated overnight at 4 °C with primary antibodies diluted in 10% goat serum and washed with Tris buffered saline (pH 7.4). For subsequent bright-field light microscopy analysis the REALTM EnVisionTM Detection System, consisting of peroxidase/diaminobenzidine+ (DAB+) rabbit/mouse (K5007, Dako, Glostrup, Denmark), was used to detect the primary antibodies. The paraffin sections were counterstained with Mayer`s hematoxylin, dehydrated in graded alcohols, and cover-slipped with Pertex mounting medium (for details on bright-field microscopy, see (38)). Additionally, the entire heads and torsos from perfusion-fixed Prox1-EGFP+ mice were decalcified over three weeks with 10% EDTA in Tris buffer (pH 7) at RT, prior to paraffin embedding and serial sectioning. Sections were processed for immunohistochemistry for EGFP protein detection as described above. A list of the primary antibodies used can be found in Table 1.

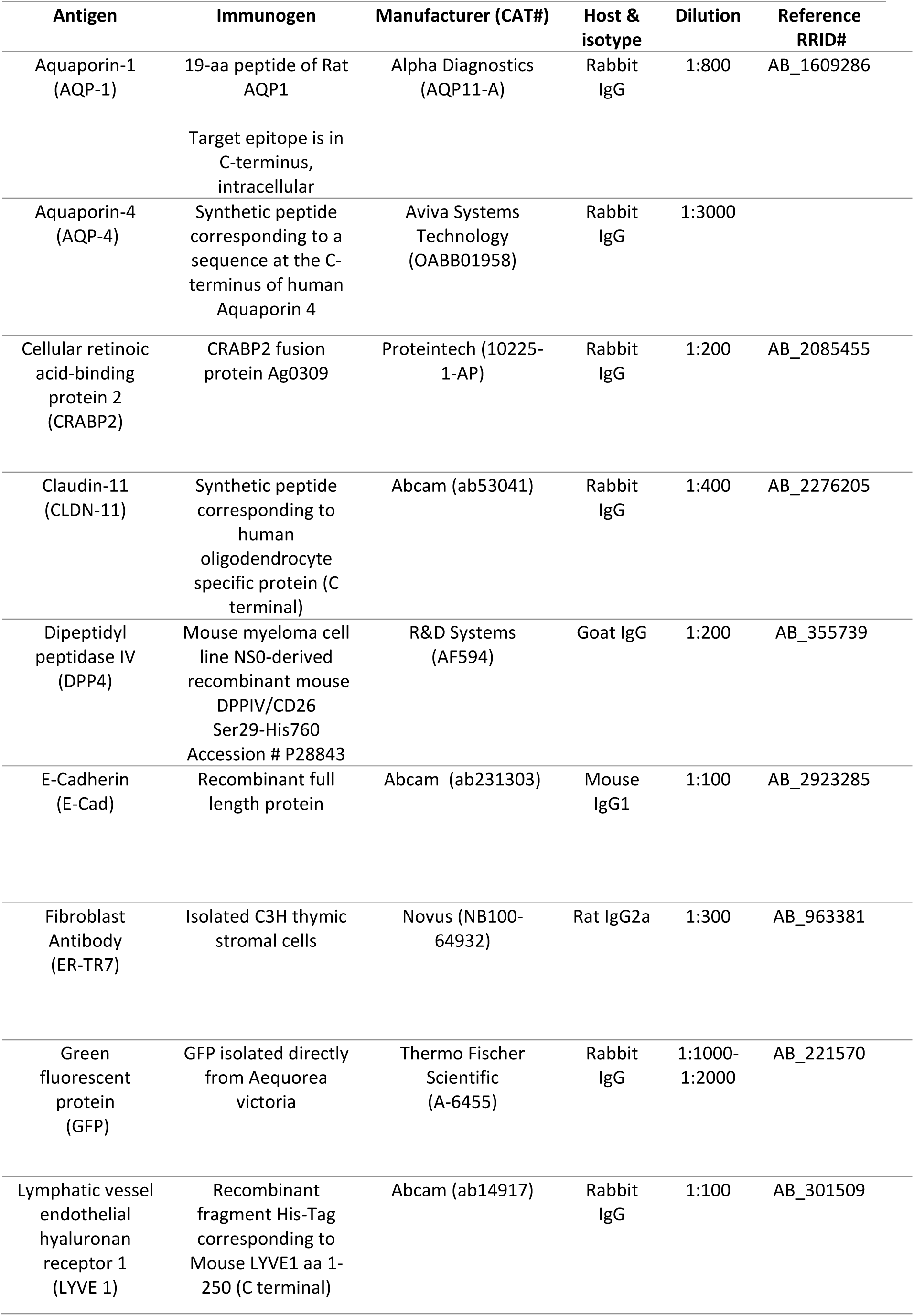
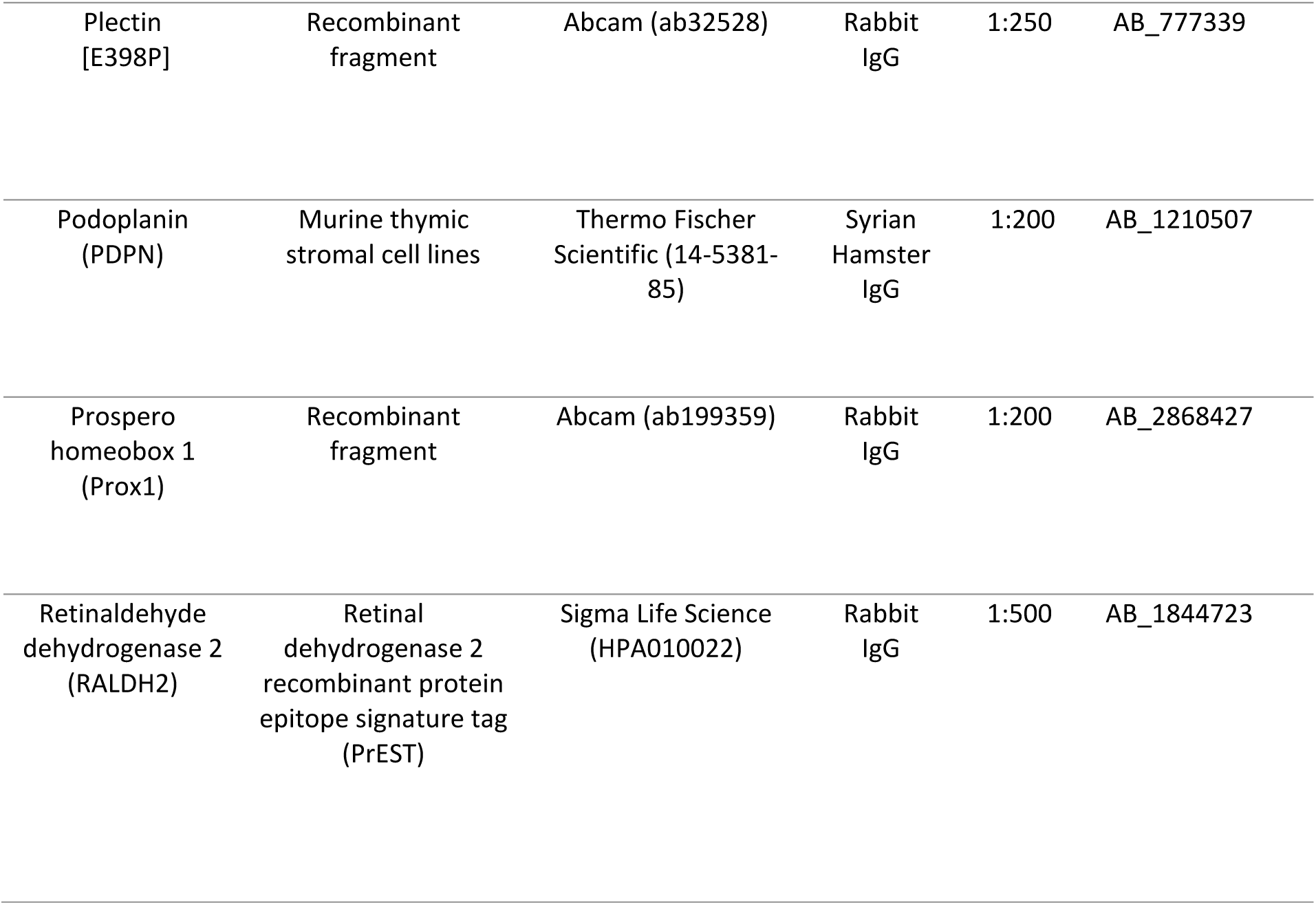

## List of abbreviations

ABC layer: Arachnoid Barrier Cell Layer
CiA: cisterna ambiens
CiC: Cisterna caroticus
CiP: cisterna pontis
CLDN-11: Claudin 11
CRABP2: Cellular retinoic acid-binding protein 2
DPP4: Dipeptidyl peptidase IV
LYVE1: Lymphatic vessel endothelial hyaluronan receptor 1
PBS: Phosphate Buffered Saline
PDPN: Podoplanin
RALDH2: Retinaldehyde dehydrogenase 2
SAS: Subarachnoid Space
SLYM: Sub-arachnoid Lymphatic-like Membrane
VEGFRC: Vascular endothelial growth factor

## Declarations

### Ethics approval and consent to participate

All experiments carried out at the University of Copenhagen were approved by the Animal Experiments Council under the Danish Ministry of Environment and Food (license number: 2015-15-0201-00535) and the procedures were performed in accordance with the European directive 2010/63/EU, with an effort was undertaken to reduce the utilization of animals. This study does not involve human participants, human data or human tissue, thus consent to participate is not applicable.

### Consent for publication

Not applicable

### Availability of data and materials

The datasets used and/or analyzed during the current study are available from the corresponding author on reasonable request.

### Authors’ contributions

KM: Conceptualization, Funding acquisition, Investigation, Methodology, Project administration, Resources, Supervision, Validation, Visualization, Writing - original draft, and Writing - review & editing.

VP: Investigation, Methodology, Project administration, Validation, Visualization, Writing - original draft, and Writing - review & editing.

MG, AL, DG: Investigation, Methodology, Visualization, and Writing - review & editing. SB: Investigation, Writing - review & editing.

MN: Conceptualization, Formal analysis, Funding acquisition, Methodology, Project administration, Resources, Supervision, Validation, Visualization, Writing - original draft, and Writing - review & editing.

YM: MRI image acquisition and analysis

## Acknowledgments

We acknowledge P. S. Froh (Department of Cellular and Molecular Medicine, Faculty of Health and Medical Sciences, University of Copenhagen, Denmark) for excellent technical assistance with the histology and immunohistochemistry of the decalcified samples. We also thank D. Xue for expert graphical support.

---

**Supplementary Fig. 1:**
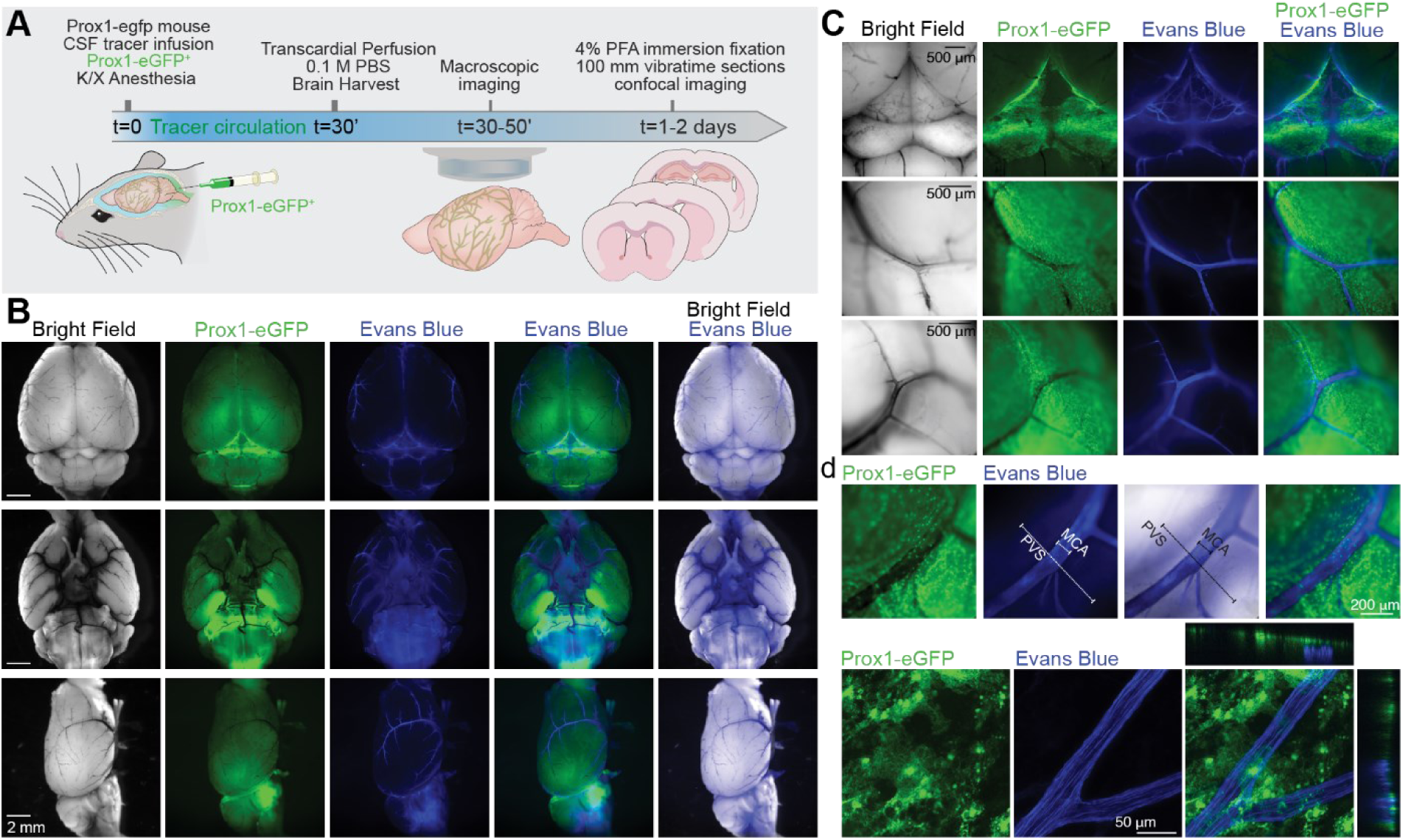
SLYM layer defines the perivascular space. **(A)** Scheme of the experimental approach. Tracer [Evans Blue, 0.9 kDa, 0.5% in aCSF (w/v), 10 µL, 2 µL/min] was delivered by cisterna magna cannulation to CSF space of K/X-anesthetized Prox1-eGFP mice. After 30 minutes circulation of the tracer, mice were perfused with PBS to eliminate blood vessel signal and brains were extracted and acutely imaged under a fluorescent microscope. **(B)** Macroscope bright field and fluorescent images of the whole brain. Bright field images (grey) show the basal cisterns filled with Evans Blue. Green signal corresponds to Prox1-eGFP expression and Evans Blue fluorescence is additionally shown in blue. **(C)** Close-up images over the pineal gland (dorsal) and MCA (ventral) illustrate the accumulation of Evans Blue in the arterial space, covered by Prox1-eGFP+ cells. **(D)** To evaluate the spatial location of Prox1-eGFP cells regarding MCA, high magnification images were combined with confocal images, where orthogonal views showed the location of Evans Blue signal on the parenchymal side below the SLYM layer.

**Supplementary Fig. 2:**
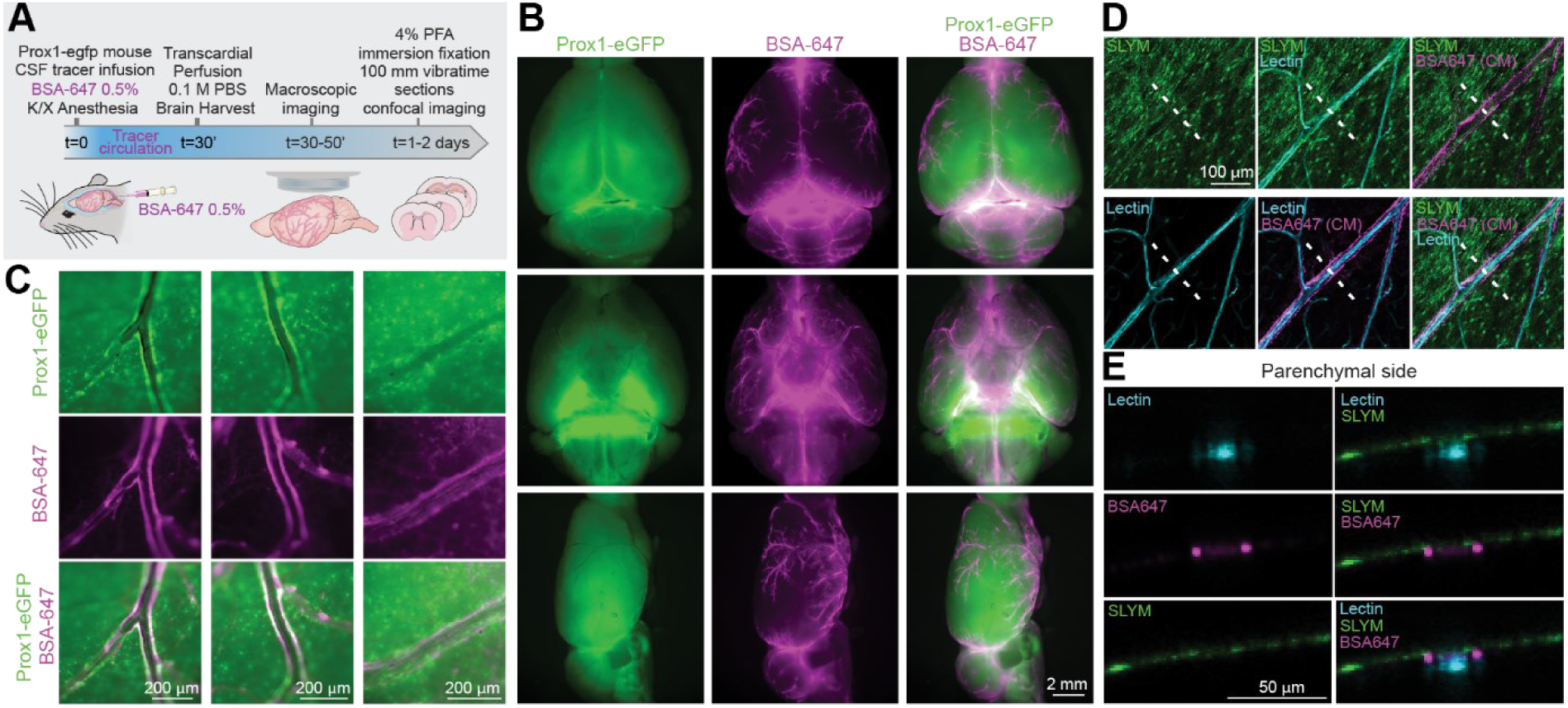
SLYM layer defines the perivascular space. **(A)** Scheme of the experimental approach. Tracer [BSA-647, 67 kDa, 0.5% (w/v), 10 µL, 2 µL/min] was delivered by cisterna magna cannulation to CSF space of K/X-anesthetized Prox1-eGFP mice. After 30 minutes circulation of the tracer, mice were perfused with PBS to eliminate blood vessel signal and brains were extracted and acutely imaged under a fluorescent microscope. **(B)** Macroscope fluorescent images of the whole brain, showing the basal cisterns filled with BSA-647 (magenta). Green signal corresponds to Prox1-eGFP expression. **(C)** Close-up images over the MCA at either the dorsal or ventral side illustrate the accumulation of BSA-647 in the periarterial space, covered by Prox1-eGFP+ cells. **(D)** Confocal Z-stacks confirmed the location of the fluorescent tracer in the perivascular space. Blood vessels were labelled internally with lectin (WGA-405, cyan) to improve the spatial location. **(E)** Reslicing of the confocal Z-stacks along the white line indicated in D showed the tridimensional distribution of the tracer (magenta) regarding SLYM (Prox1-eGFP+, green).

